# *Hnf4a* is required for the development of Cdh6-expressing progenitors into proximal tubules in the mouse kidney

**DOI:** 10.1101/2020.02.16.951731

**Authors:** Sierra S. Marable, Eunah Chung, Joo-Seop Park

## Abstract

**Background:** Hnf4a is a major regulator of renal proximal tubule (PT) development. In humans, a mutation in *HNF4A* is associated with Fanconi renotubular syndrome (FRTS), which is caused by defective PT functions. In mice, mosaic deletion of *Hnf4a* in the developing kidney causes a paucity of PT cells, leading to FRTS-like symptoms. The molecular mechanisms underlying the role of Hnf4a in PT development remain unclear.

**Methods:** We generated a new *Hnf4a* mutant mouse model employing *Osr2Cre,* which effectively deletes *Hnf4a* in developing nephrons. We characterized the mutant phenotype by immunofluorescence analysis. We performed lineage analysis to test if Cdh6-expressing cells are PT progenitors. We also performed genome-wide mapping of Hnf4a binding sites and differential gene analysis of *Hnf4a* mutant kidneys to identify direct target genes of Hnf4a.

**Results:** Deletion of *Hnf4a* with *Osr2Cre* led to the complete loss of mature PT cells, causing lethality in the *Hnf4a* mutant mice. We found that Cdh6^high^, LTL^low^ cells serve as PT progenitors and that they show higher proliferation than Cdh6^low^, LTL^high^ differentiated PT cells. We also found that Hnf4a is required for PT progenitors to develop into differentiated PT cells. Our genomic analyses revealed that Hnf4a directly regulates the expression of genes involved in transmembrane transport and metabolism.

**Conclusions:** Our findings show that Hnf4a promotes the development of PT progenitors into differentiated PT cells by regulating the expression of genes associated with reabsorption, the major function of PT cells.

**Significance:** Proximal tubule cells are the most abundant cell type in the mammalian kidney and they perform the bulk of the renal reabsorption function. Despite their importance in kidney function, the molecular mechanisms of proximal tubule development and maturation are not well understood. Here we find that, in the developing mouse kidney, Cdh6^high^, LTL^low^ cells act as proximal tubule progenitors and that Hnf4a is required for these cells to further develop into proximal tubules. Our genomic analyses show that Hnf4a directly regulate the expression of genes required for reabsorption such as transmembrane transport genes and metabolism genes. This study advances our understanding of how kidney proximal tubule cells form during development.

## INTRODUCTION

The kidneys filter the blood, regulate osmotic levels, maintain electrolyte balance, and metabolize drugs. The functional unit of the kidney is the nephron, which is composed of the glomerulus, the proximal tubule, the loop of Henle, and the distal tubule.^1^ Each segment of the nephron has distinct physiological functions and morphology. The proximal tubule cells are the most populous cell type in the kidney and they carry out the bulk of reabsorption in the nephron.^2–4^ Under physiological conditions, proximal tubules reabsorb approximately two-thirds of glomerular-filtered water and sodium chloride as well as most of the filtered glucose and phosphate.^5^ Proximal tubular reabsorption of water and metabolites is essential in the regulation of body fluid composition and volume. Numerous transporter and metabolism genes are expressed in the proximal tubules in order to facilitate the function and energy demands of these highly active renal epithelial cells.^6–12^ Despite their importance in kidney function, the molecular mechanisms of proximal tubule development and maturation are not well understood.

Fanconi renotubular syndrome (FRTS) is defined as generalized proximal tubule dysfunction.^13, 14^ Symptoms of FRTS include glucosuria, phosphaturia, proteinuria, polyuria, and polydipsia.^14, 15^ These symptoms are consistent with a failure of the proximal tubules to reabsorb and transport filtered molecules, causing urinary wasting.^16, 17^ In humans, the heterozygous mutation R76W in the *HNF4A* gene causes FRTS with nephrocalcinosis.^18^ Since this mutation is located in the DNA-binding domain, it has been speculated that the mutation affects the interactions of HNF4A with regulatory DNA.^18^ A recent study of the FRTS *HNF4A* mutation in *Drosophila* nephrocytes confirmed that the mutation reduced binding of Hnf4a to DNA and caused nuclear depletion of Hnf4a in a dominant-negative manner, leading to mitochondrial defects and lipid accumulation.^19^

We have previously shown that *Hnf4a* is expressed in developing proximal tubules in the mouse kidney and that *Hnf4a* is important for proximal tubule formation.^20^ Mosaic loss of *Hnf4a* in the murine nephron lineage caused Fanconi renotubular syndrome-like symptoms, including polyuria, polydipsia, glucosuria, and phosphaturia.^20^ Due to the mosaic expression of *Six2GFPCre* in mesenchymal nephron progenitor cells, the *Hnf4a* mutant kidney by *Six2GFPCre* was a chimera of wild-type and mutant cells.^20^ This made it difficult to perform more rigorous differential gene expression analyses. In this study, we generated a new mouse model with thorough deletion of *Hnf4a* in the proximal segments of the nephron using *Osr2^IresCre^* and investigated the requirement of mature proximal tubules for postnatal survival.^21^ We also performed lineage tracing to identify proximal tubule progenitor cells in the developing kidney. To further elucidate the role of *Hnf4a* in proximal tubule development, we performed genome-wide mapping of Hnf4a binding sites in the murine neonate kidney and transcriptomic analysis of the *Hnf4a* mutant kidney. We found that Hnf4a is required for terminal differentiation of proximal tubule cells and that mature proximal tubules are required for postnatal survival. Cdh6^high^, LTL^low^ cells in the developing kidney are proximal tubule progenitor cells and loss of *Hnf4a* causes their developmental arrest. Our genomic analyses revealed that Hnf4a directly regulates expression of many mature proximal tubule genes, including transport and metabolism genes, consistent with the fact that active reabsorption is the major function of proximal tubules.

## METHODS

### Mice

All mouse alleles used in this study have been previously published: *Qsr2^tm2(cre)Jian^* (*Osr2^IresCre^*);^22^ *Hnf4a^tm1Sad^* (*Hnf4a^c^*);^23^ *Cdh6^tm1.1(cre/ERT2)Jrs^* (*Cdh6^CreER^*);^24^ *Gt(ROSA)26Sor^tm3(CAG-EYFP)Hze^* (*Rosa26^Ai3^*).^25^ All mice were maintained in the Cincinnati Children’s Hospital Medical Center (CCHMC) animal facility according to animal care regulations. All experiments were performed in accordance with animal care guidelines and the protocol was approved by the Institutional Animal Care and Use Committee of the Cincinnati Children’s Hospital Medical Center (IACUC2017-0037). We adhere to the NIH Guide for the Care and Use of Laboratory Animals.

### Tamoxifen Treatment

Tamoxifen (T5648, Sigma) was dissolved in corn oil (C8267, Sigma) at a concentration of 20mg/ml. Pregnant female mice were injected with tamoxifen intraperitoneally (4mg/40g body weight).

### Immunofluorescence Staining

Embryonic, neonatal, and adult murine kidneys were fixed in 4% paraformaldehyde in phosphate-buffered saline (PBS), incubated overnight in 10% sucrose/PBS at 4°C, and imbedded in OCT (Fisher Scientific). Cryosections (8-9μm) were incubated overnight with primary antibodies in 5% heat-inactivated sheep serum/PBST (PBS with 0.1% Triton X-100). We used primary antibodies for GFP (1:500, Aves GFP-1020), Jag1 (1:20, DSHB TS1.15H), Wt1 (1:100, Santa Cruz sc-7385), Biotin-LTL (1:500, Vector Labs B-1325), FITC-LTL (1:200, Vector Labs FL-1321), Hnf4a (1:500, Abcam ab41898), Hnf4a (1:500, Santa Cruz sc-8987), Lrp2 (1:100, Santa Cruz sc-515772), Ki67 (1:500, BioLegend 652402), Slc12a1 (1:500, Proteintech 18970-1-AP), Slc12a3 (1:300, Sigma HPA028748), Ass1 (1:500, Proteintech 16210-1-AP), Slc5a2 (1:100, Sigma HPA041603), Miox (1:500, Sigma HPA039562), Cdh6 (1:100, R&D Systems MAB2715-SP), and Cdh6 (1:200, Sigma HPA007047). Fluorophore-labeled secondary antibodies were used for indirect visualization of the target. Images were taken with a Nikon Ti-E widefield microscope equipped with an Andor Zyla camera and Lumencor SpectraX light source housed at the Confocal Imaging Core (CIC) at CCHMC.

### Histology

Mouse kidneys were harvested and fixed in 4% paraformaldehyde in PBS overnight. Paraffin sections (5 μm) were stained with hematoxylin and eosin or periodic acid-Schiff reagent (American MasterTech KTPAS). Images were taken with a Nikon Ti-E widefield microscope equipped with an Andor Zyla camera and Lumencor SpectraX light source housed at the Confocal Imaging Core at CCHMC.

### Cell counts

The quantification of Ki67 staining was performed on *Hnf4a^c/c^* and *Hnf4a^c/+^;Osr2^IresCre^* P0 control kidneys. Immunostained samples were imaged as described above. Three 250,000μm^2^ sections from each pair of kidneys (four pairs of kidneys total) were quantified manually using the ImageJ multi-point tool.^26^ For the quantification of Cdh6^+^, LTL^+^ cells, immunostained sections from *Hnf4a^c/+^;Osr2^IresCre^* (control) and *Hnf4a^c/c^;Osr2^IresCre^* (mutant) P0 kidneys were analyzed. Three 250,000um^2^ sections from each pair of kidneys (three pairs of kidneys from each genotype) were quantified manually using ImageJ multi-point tool.^26^

### ChIP-seq

ChIP-seq was performed as previously described.^27^ Briefly, kidneys from newborn (P0) mice were crosslinked with 1% paraformaldehyde, sonicated, incubated with Hnf4a antibody (Abcam ab41898) coupled Dynabeads Protein G (ThermoFisher). Non-immunoprecipitated chromatin (starting material) was used as the control. Eluted DNA was used for constructing sequencing libraries using the ThruPLEX DNA-seq kit (Takara). Libraries were sequenced on an Illumina HiSeq 2500 by the DNA Sequencing and Genotyping Core at CCHMC. ChIP-seq reads were mapped to mm9 using Bowtie.^28^ We performed peak calling and motif analysis using HOMER.^29^ Data are available at Gene Expression Omnibus under accession number GSE144824.

### RNA-seq

RNA-seq was performed as previously described.^20^ Briefly, 1 μg total RNA was isolated from P0 *Hnf4a* mutant and control kidneys using the RNeasy Plus Micro kit (Qiagen 74034) followed by mRNA was isolation with NEBNext Poly(A) mRNA Magnetic Isolation Module (E7490, NEB). Fragmentation of mRNA followed by reverse transcription and second strand cDNA synthesis was done using NEBNext Ultra RNA Library Prep Kit for Illumina (E7530, NEB). Sequencing libraries were constructed using ThruPLEX DNA-seq kit (Takara R400428). Sequencing was performed as described above. RNA-seq reads were mapped to mm9 using TopHat and normalized gene expression values were calculated using Cufflinks.^30^ Genes that showed at least a 1.5-fold change in expression with a *p*-value ≤0.05 were considered differentially expressed. Data are available at Gene Expression Omnibus under accession number GSE144772.

### Genomic Regions Enrichment of Annotations Tool (GREAT) analysis

GREAT analysis was performed using the online program, version 3 (great.stanford.edu).^31^ To associate genomic regions with genes, gene regulatory domains were defined as minimum 5.0 kb upstream and 1.0 kb downstream of the TSS, and distally up to 1000 kb to the nearest gene’s basal domain (‘basal plus extension’ option). 10,417 genomic regions from the Hnf4a ChIP-seq dataset were entered into the GREAT online program and Mouse Genome Informatics (MGI) Expression terms of genes associated with the genomic regions were assessed.

### Gene Ontology Analysis

Gene ontology analysis was performed using DAVID Bioinformatics Resources (david.ncifcrf.gov) on differentially expressed genes identified from the RNA-seq analysis.^32^

### Statistical Analyses

Statistical analysis of Kaplan-Meier survival curve was performed using GraphPad 8 Prism software (*n*=11 *Hnf4a* mutants and *n*=14 controls).^33^ The Log-rank test was used for survival analysis. Student’s *t* test was performed using GraphPad 8 Prism software. P<0.05 was considered to be significant.

## RESULTS

### Hnf4a is required for mature proximal tubule formation

In our previous study, we utilized *Hnf4a* floxed alleles and *Six2GFPCre* to generate a mouse model with kidney-specific deletion of *Hnf4a.* However, *Six2GFPCre* displayed mosaic expression in nephron progenitor cells and allowed a subset of nephron progenitors to escape Cre-mediated recombination.^20, 34, 35^ Therefore, our previous *Hnf4a* mutant kidney was a chimera of wild-type and mutant cells leading to the FRTS-like phenotype we observed. In order to thoroughly investigate the *Hnf4a* loss-of-function phenotype, we utilized a less mosaic Cre that specifically targets the proximal segments of the nephron. We generated a new mouse model with nephron-specific deletion of *Hnf4a* using a mouse line expressing Cre recombinase under the *Osr2* promoter *(Osr2^IresCre^)^22^* bred with *Hnf4a* floxed mice (*Hnf4a^c/c^*).^23^ We have recently shown that *Osr2^IresCre^* is expressed in the proximal and medial segments of the S-shaped body (SSB) of the developing nephron and that the medial segment of SSB develops into proximal tubules and loops of Henle.^21^ This Cre, therefore, targets all nephron segments except for the distal tubule. *Hnf4a* deletion was apparent as early as the SSB stage in the *Hnf4a* mutant kidney, and loss of *Hnf4a* did not appear to affect SSB formation (Supplemental Figure 1). *Osr2^IresCre^* achieved almost complete deletion of *Hnf4a* in the kidney (Figure 1A). *Lotus Tetragonolobus* Lectin (LTL) is known to bind to glycoproteins on the surface of the proximal tubules specifically. Deletion of *Hnf4a* in the nephron led to the loss of differentiated proximal tubule cells with high LTL staining (LTL^high^) in postnatal day 0 (P0) kidneys (Figure 1A) but it did not affect the formation of other nephron segments (Supplemental Figure 2A). Consistent with this, our transcriptomic analysis also showed that only proximal tubule genes were significantly downregulated in the *Hnf4a* mutant (Supplemental Figure 2B).

**Figure 1.**
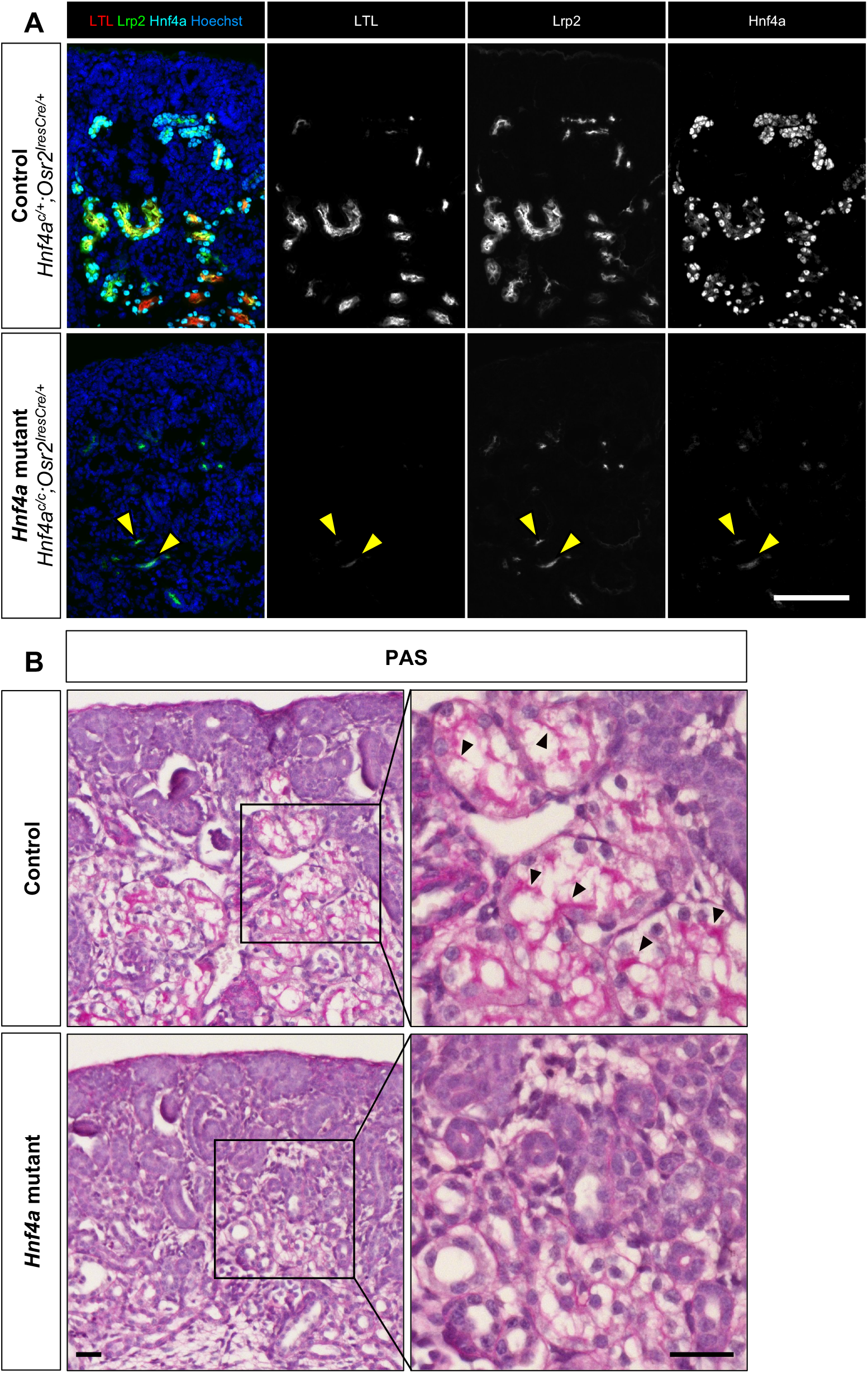
*Hnf4a* deletion by *Osr2Cre* leads to loss of mature proximal tubule (PT) cells. (A) Loss of *Hnf4a* in the nephron inhibits formation of LTL^high^, mature PT cells and causes a decrease in expression of *Lrp2,* a PT-specific gene in the newborn (P0) kidney. Yellow arrowheads mark LTL^low^, Lrp2^low^ cells that persist in the mutant. Image is representative of *n*=3. Scale bar, 100μm. (B) Periodic acid-Schiff (PAS) staining of control and *Hnf4a* mutant kidneys at P0 show *Hnf4a* mutants lack brush border. Black arrowheads mark brush border. Image is representative of *n*=3. Scale bar, 50μm.

The *Hnf4a* mutant kidneys showed a decrease in the level of Lrp2 (Low-density lipoprotein-related protein 2), a proximal tubule-specific endocytic receptor protein also known as Megalin (Figure 1A).^20, 38^ A few LTL^low^, Lrp2^low^ cells persisted in the *Hnf4a* mutant kidney (Figure 1A, yellow arrowheads). We reasoned that these cells might represent immature proximal tubules or proximal tubule progenitor cells and that the *Hnf4a* mutant kidney lacks mature proximal tubules. From previously published single cell RNA-seq (scRNA-seq) data,^8^ we identified *Ass1, Slc5a2,* and *Miox* as genes that are highly expressed in mature proximal tubules and examined their expression in the *Hnf4a* mutant kidney. We found that the *Hnf4a* mutant kidney showed little to no staining of Slc5a2, Ass1, and Miox, indicating decreased expression of mature proximal tubule genes in the *Hnf4a* mutant kidney (Supplemental Figure 3). A distinctive feature of the mature proximal tubules is the apical brush border. The brush border is composed of microvilli which increase the surface area of the proximal tubule to facilitate reabsorption.^9, 10, 39^ The absence of the proximal tubule brush border has been associated with proximal tubule dysfunction in patients, highlighting the importance of brush border formation.^40^ We analyzed brush border formation using the Periodic acid-Schiff (PAS) stain^41^ and found that *Hnf4a* mutants showed a lack of brush border formation (Figure 1B). Considering that brush border formation only occurs in postmitotic, differentiated cells,^42^ our result suggests that the *Hnf4a* mutant kidney lacks terminally differentiated proximal tubule cells.

Due to *Hnf4a* mutant kidneys lacking mature proximal tubules, we examined whether the nephron tubules were still connected to the glomerulus in the mutant kidney. Immunofluorescence staining showed that the glomerulus was connected to Cdh6^+^ cells in both the control and *Hnf4a* mutant kidneys but that LTL staining was visible only in the control kidney (Supplemental Figure 4), suggesting that the glomerulus is conncected to proximal tubule progenitors in the *Hnf4a* mutant kidney.

### Loss of *Hnf4a* in the nephron leads to postnatal lethality

To examine the effects of loss of LTL^high^ differentiated proximal tubules on postnatal kidney development, we analyzed the histology of *Hnf4a* mutant kidneys at P0, P7, and P14. At P0, the *Hnf4a* mutant kidney was similar in size to the control (Figure 2A). At P7, the *Hnf4a* mutant kidney was slightly smaller with a thinner cortex than the control kidney (Figure 2B). There was no apparent defect in glomerulus, loop of Henle, or distal tubule formation in the *Hnf4a* mutant at P7 (Supplemental Figure 5). At P14, the medullary region of the *Hnf4a* mutant kidney was severely damaged, cysts formed in the cortical region, and hydronephrosis was apparent (Figure 2C). Increased filtrate flow through the renal tubules can lead to renal pelvic dilation and nonobstructive hydronephrosis in nephrogenic diabetes insipidus.^43–45^ It is likely that hydronephrosis seen in the *Hnf4a* mutant kidney is caused by increased filtrate flow through the nephron tubules due to lack of reabsorption in the proximal tubule. Survival analysis of the *Hnf4a* mutant mice showed that ~60% of *Hnf4a* mutants were deceased by P14, likely due to kidney dysfunction (Figure 2D). No *Hnf4a* mutants survived to weaning age (P28). These results show that the lack of mature proximal tubules causes postnatal lethality, highlighting the importance of the mature proximal tubule function for survival.

**Figure 2.**
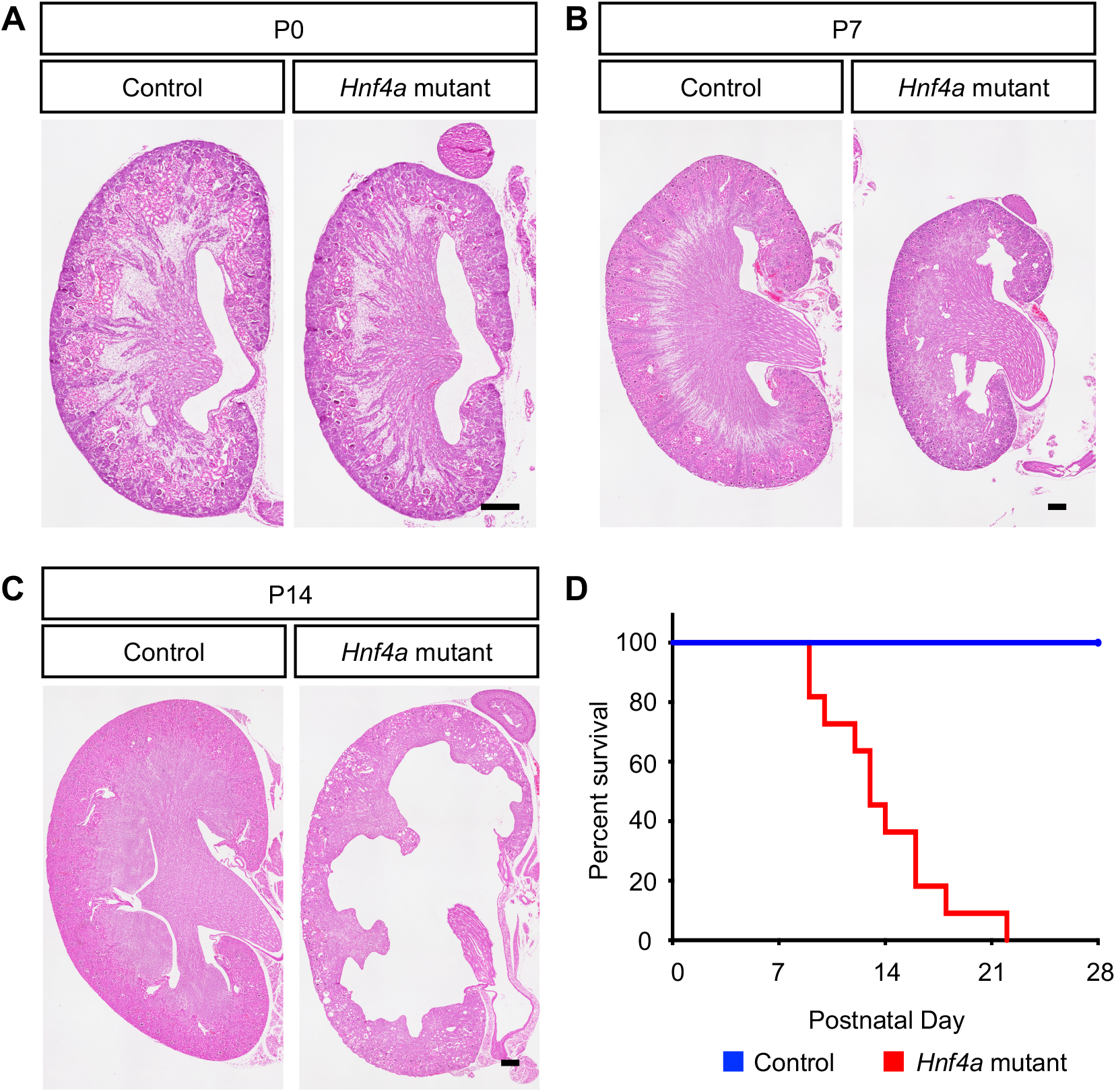
Loss of mature proximal tubules leads to postnatal lethality in *Hnf4a* mutant mice. (A-C) Hematoxylin and eosin (H&E) staining of *Hnf4a* mutant kidneys at birth (P0), postnatal day 7 (P7), and postnatal day 14 (P14). Images are representative of *n*=4. Scale bar, 100μm. (D) Kaplan-Meier survival analysis of the *Hnf4a* mutants (*n*=11) with heterozygous controls (*n*=14). *P-value < 0.0001, determined by Log-rank test.

### Cdh6^high^, LTL^low^ cells in the developing kidney are proximal tubule progenitor cells

It has been previously suggested that Cdh6-expressing cells in the developing murine kidney are presumptive proximal tubule cells.^46^ It was reported that, in the mouse embryonic kidney, *Cdh6* was expressed in the medial segment of the SSB and that LTL^+^ proximal tubules were still positive for Cdh6 although its expression was downregulated compared to Cdh6^+^ cells in the nephrogenic zone. Based on these observations, it was proposed that Cdh6-expressing cells in the nephrogenic zone were destined to become proximal tubules.^46^ Consistent with this, we found that there were two distinct populations of Cdh6^+^ cells in the wild type developing murine kidney: Cdh6^high^ and Cdh6^low^ cells (Figure 3A). The majority of Cdh6^high^ cells were Hnf4a^+^ and had no or low LTL staining (red arrowheads and orange arrowheads, respectively, in Figure 3A), suggesting that these Cdh6^high^, LTL^low^ cells are prospective, immature proximal tubule cells. Cdh6^low^ cells were also positive for Hnf4a and had strong LTL staining (yellow arrowheads in Figure 3A), suggesting that these Cdh6^low^, LTL^high^ cells are differentiated proximal tubule cells. We found that, in the *Hnf4a* mutant kidney, Cdh6^low^, LTL^high^ cells were absent and the number of Cdh6^high^, LTL^low^ cells were increased, suggesting that the loss of Hnf4a prevents Cdh6^high^, LTL^low^ cells from developing into Cdh6^low^, LTL^high^ cells (Figure 3B). We found that *Cdh6* expression in the kidneys persisted postnatally, suggesting continued proximal tubule development and maturation (Supplemental Figure 6).

**Figure 3.**
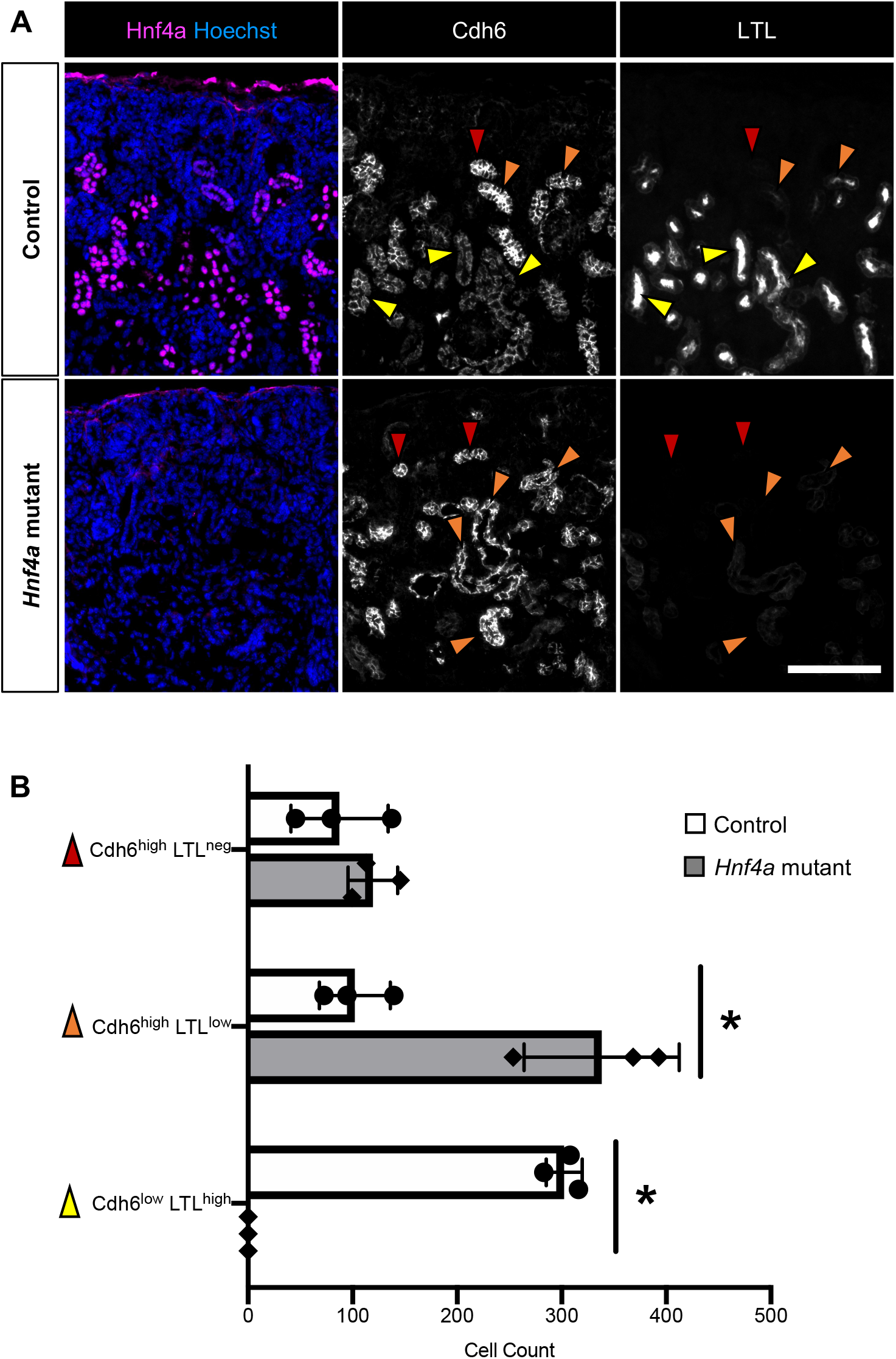
High Cdh6 expression is persistent in the *Hnf4a* mutant kidney. (A) In the P0 control kidney, Cdh6 expression is high in LTL^neg^ and LTL^low^, presumptive PT cells (red and orange arrowheads, respectively) and Cdh6 expression decreases as PT cells develop into LTL^high^, mature PT cells (yellow arrowheads). In the *Hnf4a* mutant kidney, Cdh6^high^,LTL^low^ cells are more abundant compared to the control and there are no Cdh6^low^, LTL^high^ cells to be found. Scale bar, 100μm. Image is representative of *n*=3. (B) Quantification of Cdh6^high^ and Cdh6^low^ cells in the *Hnf4a* mutant and control kidney. *P-value < 0.01, determined by *t* test.

In order to definitively test if Cdh6^high^ cells are proximal tubule progenitor cells, we performed lineage analysis using a tamoxifen-inducible *Cre* recombinase under the *Cdh6* promoter *(Cdh6^CreER^)* and a Cre-inducible *Rosa26^Ai3^* reporter.^24, 25^ If Cre-mediated activation of the Rosa reporter is restricted to the proximal tubules and not detected in other nephron segments, it would suggest that Cdh6+ cells are proximal tubule progenitors. Pregnant dams were injected with tamoxifen at E14.5 or E16.5 to label Cdh6^high^ cells and their descendant cells with the *Rosa26^Ai3^* reporter. Embryos were harvested at E18.5. We found that all *Rosa26^Ai3^* labeled cells were Hnf4a^+^ and most were also LTL^+^. As expected, transient activation of Cre acitivity by tamoxifen was not able to produce a robust labeling of proximal tubules with the Rosa reporter, partly due to the fact that the formation of nephrons is an asynchronous process in the rapidly developing kidney. Nonetheless, our observation that the descendants of Cdh6+ cells differentiate into proximal tubules *exclusively* indicates that Cdh6^high^ cells in the developing kidney are proximal tubule progenitor cells (Figure 4).

**Figure 4.**
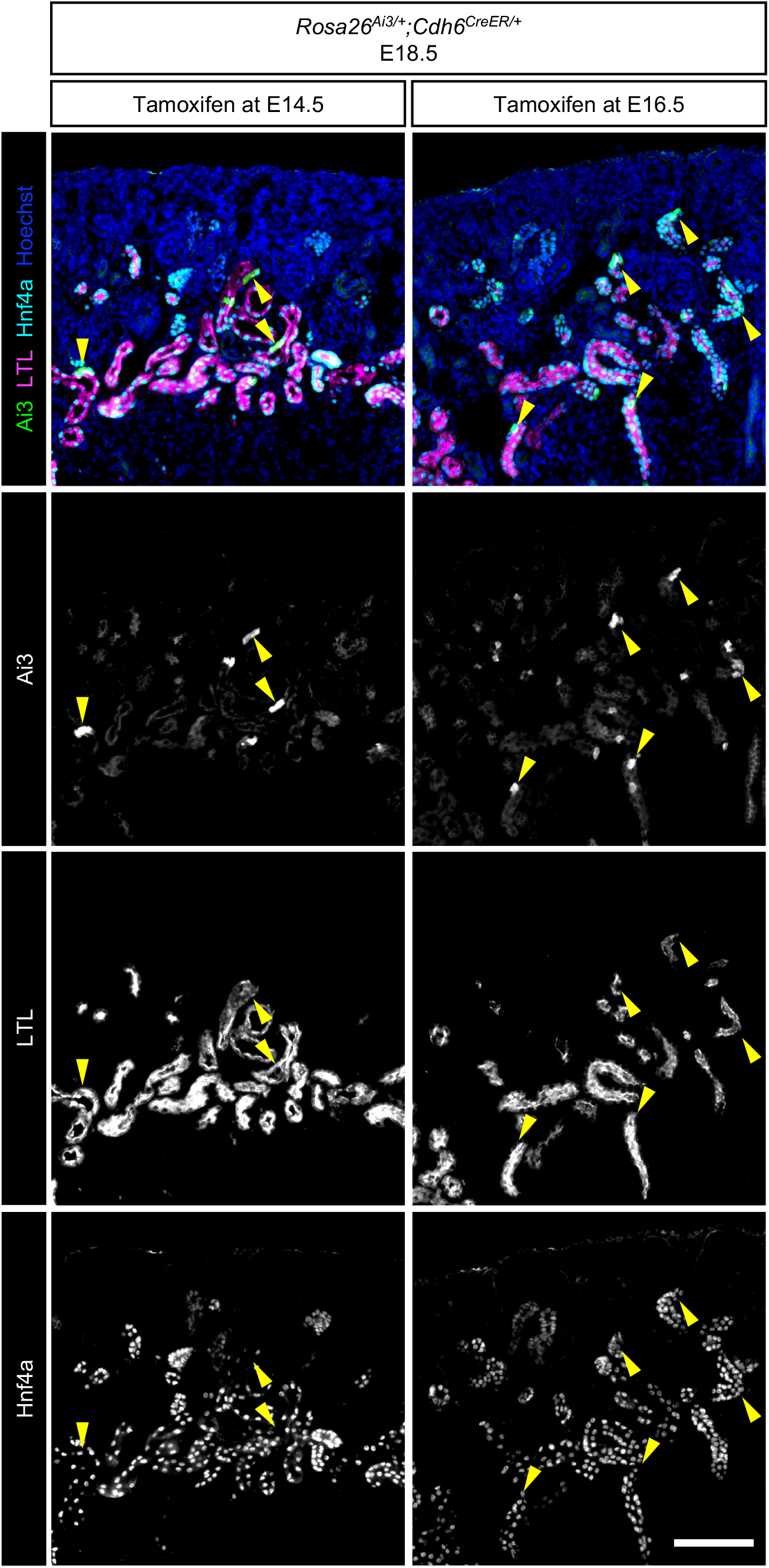
Cdh6 lineage tracing shows that Cdh6^+^ cells are PT progenitor cells. Lineage labeling of Cdh6^+^ cells with Ai3 after tamoxifen injection into pregnant dams at E14.5 or E16.5. All Ai3^+^ cells are Hnf4a^+^ and most are also LTL^+^ (yellow arrowheads) at E18.5. Images are representative of *n*=3. Scale bar, 100μm.

Hnf4a has been shown to inhibit proliferation in hepatocytes and promote terminal differentiation.^47^ Many models of cellular differentiation show an inverse relationship between proliferation and differentiation.^48–51^ Terminal differentiation commonly involves exiting the cell cycle and entering a postmitotic state.^48, 52^ To determine whether the transition from Cdh6^high^, LTL^low^ proximal tubule progenitors to Cdh6^low^, LTL^high^ differentiated proximal tubule cells coincides with cell cycle exit, we examined Ki67 expression in Cdh6^high^ and Cdh6^low^ cell populations (Figure 5, A and B).

**Figure 5.**
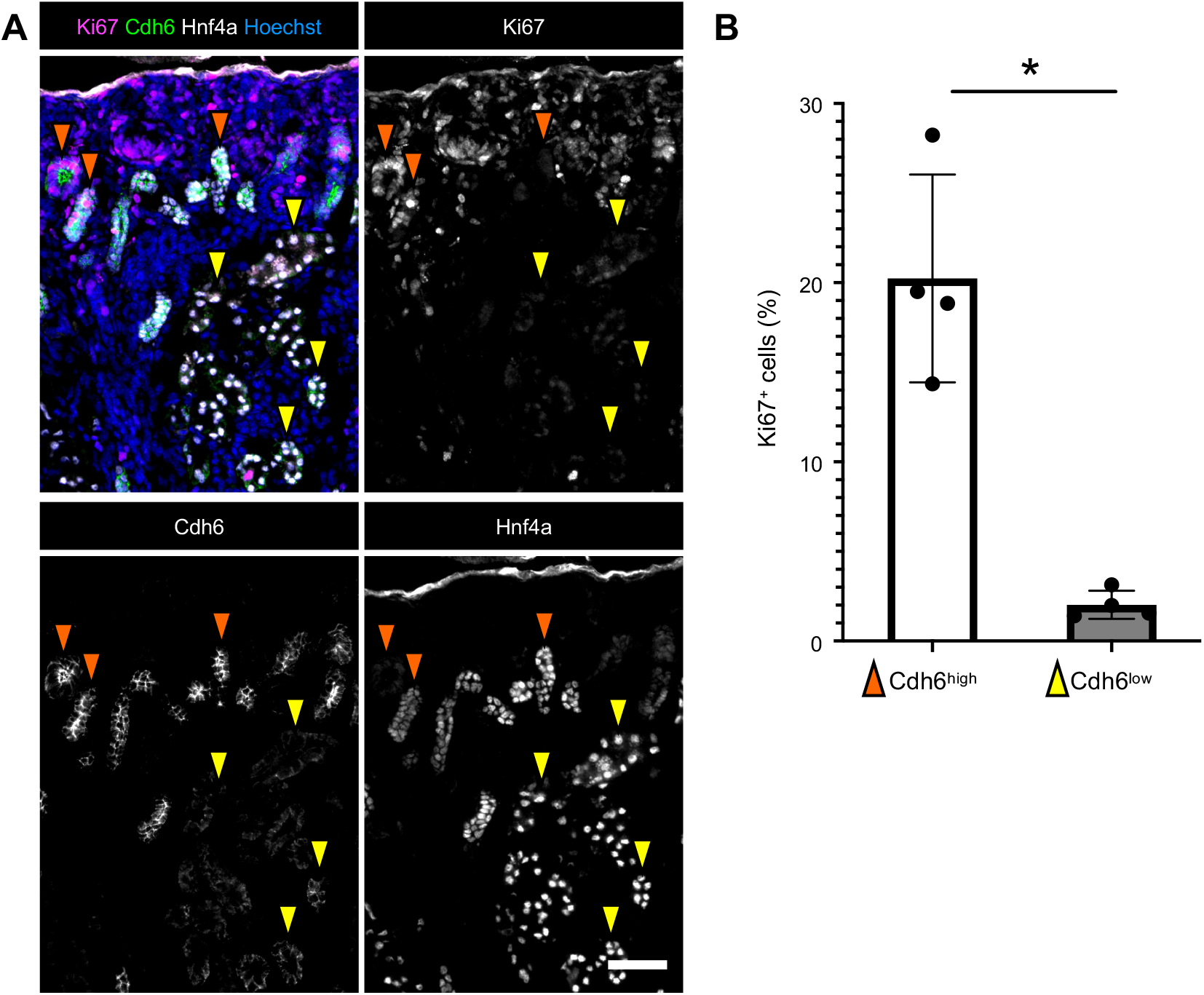
Cdh6^high^ PT progenitor cells have a higher proliferation rate than Cdh6^low^ mature PT cells. (A) Representative immunostains for Ki67, Cdh6, and Hnf4a in the P0 kidney *(n*=4). Cdh6^high^ (orange arrowheads); Cdh6^low^ (yellow arrowheads). Scale bar, 50μm. (B) Quantification of Ki67 positive cells *(n*=4). *P-value < 0.01, determined by *t* test.

Ki67 is present in actively proliferating cells and absent in resting cells.^53–56^ We found that Cdh6^high^ proximal tubule progenitor cells were highly proliferative while only few Cdh6^low^ cells showed Ki67 expression (Figure 5A). When quantified, Cdh6^high^ cells had a 10-fold higher proliferative rate than Cdh6^low^ cells, indicating an expansion of the progenitor cell population before they exit the cell cycle and undergo terminal differentiation into mature proximal tubule cells (Figure 5B). This suggests that the number of proximal tubule cells in the newborn kidney is largely determined by the proliferation of Cdh6^high^ proximal tubule progenitor cells.

### Hnf4a gene regulatory network reveals the roles of Hnf4a in regulating proximal tubule development

To further elucidate the role of Hnf4a in the proximal tubule transcriptional program, we performed chromatin immunoprecipitation followed by high-throughput sequencing (ChIP-seq) with P0 murine kidneys to identify Hnf4a-bound genomic regions. Two Hnf4a ChIP-seq replicates yielded 10,417 reproducible binding sites (peaks) (Supplemental Table 1). Our motif analysis showed high enrichment of the canonical Hnf4a binding DNA sequence within these Hnf4a peaks (Figure 6A), indicating that Hnf4a-bound genomic regions were successfully enriched in our ChIP-seq samples. We also found that DNA motifs for other nuclear receptors including the estrogen-related receptor alpha (ESRRA), the retinoid X receptor (RXR), the peroxisome proliferator-activated receptor (PPAR), and hepatocyte nuclear factor 1 beta (HNF1B) were also enriched within the Hnf4a-bound genomic regions, suggesting that these nuclear receptors share common target genes with Hnf4a to regulate proximal tubule development (Figure 6A). The majority of the Hnf4a peaks were found within 50kb of transcription start sites (TSS) (Figure 6B) and 30% of the peaks were located in promoter regions (Figure 6C). Genomic Regions Enrichment of Annotations Tool (GREAT) analysis of MGI expression annotations of genes associated with Hnf4a binding sites showed enrichment within the developing renal proximal tubules (Figure 6D),^31^ consistent with the fact that *Hnf4a* is specifically expressed in proximal tubules in the kidney. Multiple peaks were identified near the promoters of proximal tubule genes (Supplemental Table 1). In particular, we found Hnf4a peaks near genes such as *Slc34a1* and *Ehhadh,* genes linked to FRTS in human patients (Figure 6E).^57, 58^

**Figure 6.**
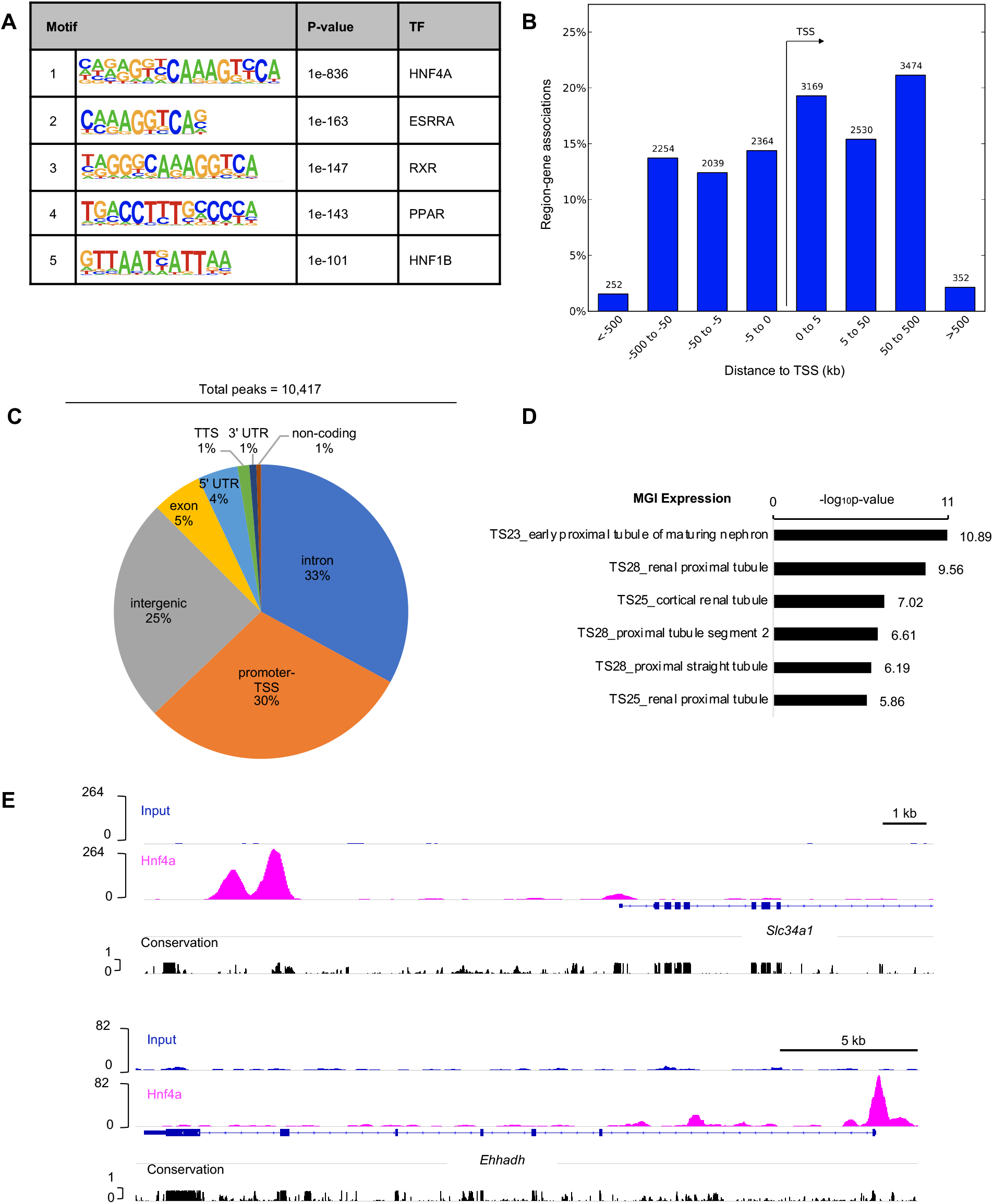
Genome-wide mapping of Hnf4a binding sites in the newborn mouse kidney. (A) Analysis of known motifs from Hnf4a ChIP-seq. (B) Bar graph showing the percentage of region-gene associations according to genomic regions’ distance to TSS computed by Genomic Regions Enrichment of Annotations Tool (GREAT) (http://bejerano.stanford.edu/great/public/html/). (C) Pie chart representing the distribution of Hnf4a peaks within the annotated genome. (D) GREAT MGI Expression annotations of Hnf4a peaks showing top six enriched terms. (E) Genome browser view of Hnf4a ChIP-seq peaks near TSS of PT genes.

We conducted transcriptomic analysis (RNA-seq) of P0 *Hnf4a* mutant kidneys to complement our candidate target gene list with differential gene expression data (Supplemental Table 2). In the *Hnf4a* mutant kidney, 442 genes showed a significant decrease in expression (fold change ≥ 1.5; p-value < 0.05), including *Slc34a1, Ehhadh,* and *Ass1*, which are highly expressed in proximal tubules (Figure 7A, Supplemental Table 2).^7, 8^ As previously mentioned, mutations in *SLC34A1* and *EHHADH* are associated with FRTS in patients.^57, 58^ Gene ontology (GO) analysis of these 442 downregulated genes showed enrichment of genes associated with metabolism and transport (Figure 7B). We found that 196 genes were significantly upregulated in the *Hnf4a* mutant kidneys (fold change ≥ 1.5; p-value < 0.05), including *Cdh6,* the gene marking proximal tubule progenitors (Figure 7A, Supplemental Table 2). GO analysis of the 196 upregulated genes showed enrichment of genes associated with phospholipid homeostasis and cholesterol transport (Figure 7C). In order to determine which genes are directly regulated by Hnf4a, we compared the differentially expressed genes in the *Hnf4a* mutant with the 7,823 genes associated with Hnf4a binding sites to find overlapping genes (Figure 7D, Supplemental Table 3). There were 245 genes in common between the significantly downregulated genes and the Hnf4a binding sites, and 81 common genes between significantly upregulated genes and the Hnf4a binding sites (Figure 7D, Supplemental Table 3). GO analysis of the 245 downregulated genes showed enrichment of genes associated with transport and metabolism (Figure 7E). Of the downregulated genes, Hnf4a has previously been shown to activate the expression of *Lrp2, Pdzk1, Slc22a1,* and *Pck1 in vitro,* a validation that some of the downregulated genes from our data are directly regulated by Hnf4a.^59–62^ When we compared our 245 downregulated genes with proximal tubule genes identified from previously published scRNA-seq,^8^ we found that there were 149 genes in common. These 149 genes were expressed mainly in mature proximal tubule populations rather than in progenitor cells (Supplemental Figure 7), implying that Hnf4a may be more active in mature proximal tubules. GO analysis of the 81 upregulated genes showed enrichment of genes associated with transport, metabolism, and response to thyroid hormone and ischemia (Figure 7F). Our genomic and transcriptomic analyses suggest that Hnf4a regulates proximal tubule maturation via activation of transport and metabolism genes, consistent with the functions of the proximal tubule.

**Figure 7.**
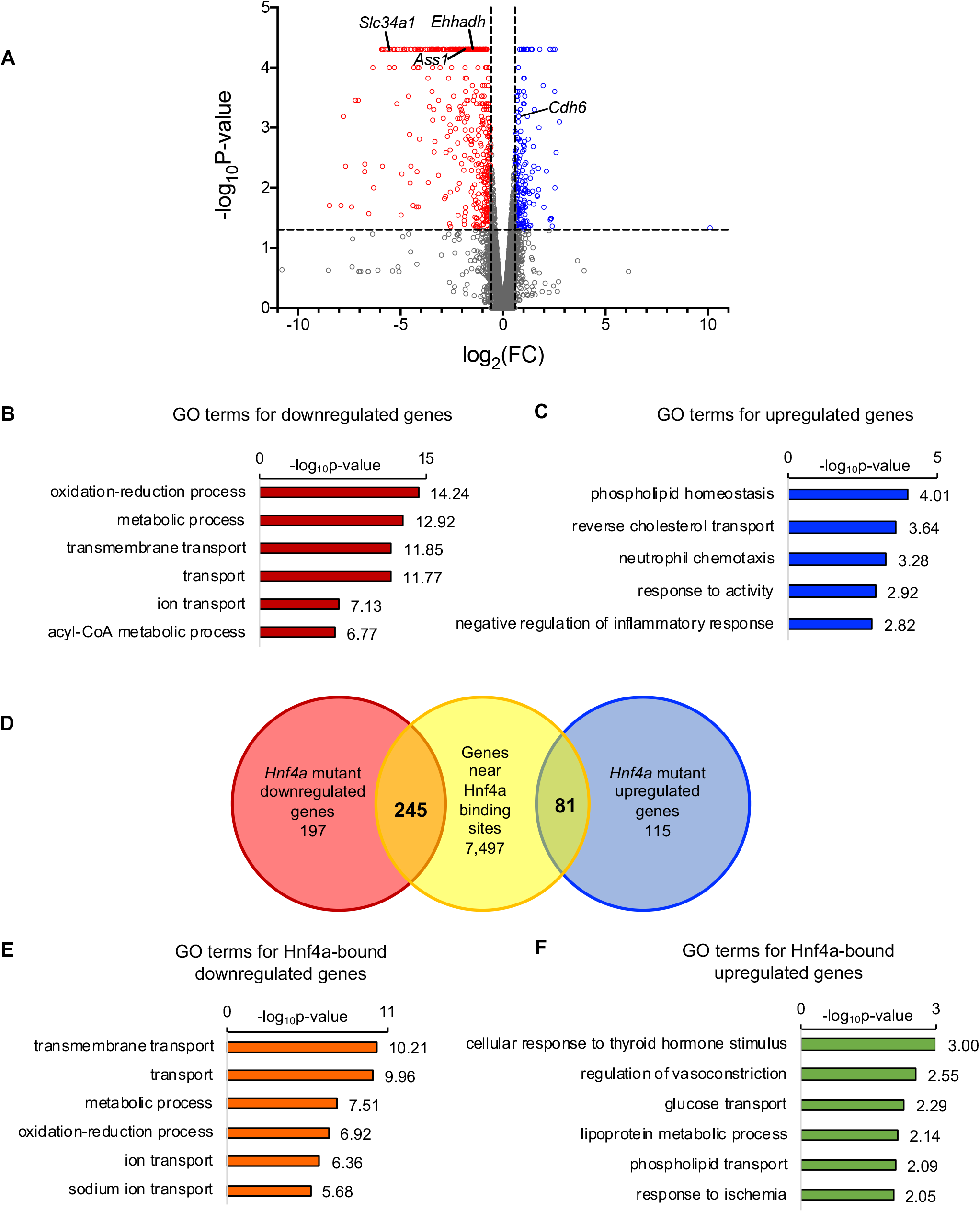
Intersection of Hnf4a ChIP-seq peaks with differentially expressed genes in the *Hnf4a* mutant kidney identified direct target genes of Hnf4a. (A) Differential expression analysis in the *Hnf4a* mutant versus the *Hnf4a* control kidney at P0. Red and blue points in the volcano plot mark genes with significantly decreased or increased expression, respectively, in the *Hnf4a* mutant. Vertical dash lines (x-axis) mark log_2_(1.5). Horizontal dash line (y-axis) marks -log_10_(0.05) (B) Gene ontology (GO) analysis of significantly downregulated genes in the *Hnf4a* mutant kidney showing top six enriched terms. (C) GO analysis of significantly upregulated genes in the *Hnf4a* mutant kidney showing top five enriched terms. (D) Venn diagram shows the overlap of genes associated with Hnf4a binding sites and differentially expressed genes in the Hnf4a mutant kidney. (E) GO analysis of Hnf4a-bound, downregulated genes showing top six enriched terms. (F) GO analysis of Hnf4a-bound, upregulated genes showing top six enriched terms.

## DISCUSSION

In our previous study, we identified two populations of LTL^+^ cells in the developing mouse kidney: LTL^low^ and LTL^high^. Based on our results, we concluded that these two populations represent presumptive proximal tubules and differentiated proximal tubules, respectively.^20^ In order to further examine the presumptive proximal tubule population, we sought to identify a marker of proximal tubule progenitors. Previously, Cdh6 had been proposed as a marker for prospective proximal tubule cells.^46^ However, it had not been definitively shown that Cdh6^+^ cells develop into proximal tubule cells.^46^ To address this, we performed lineage analysis of Cdh6^+^ cells in the developing kidney and found that these cells all became Hnf4a^+^ proximal tubule cells (Figure 4). This experiment provides strong evidence that Cdh6^+^ cells are proximal tubule progenitors in the developing kidney. Cdh6^+^ proximal tubule progenitors are highly proliferative, while LTL^high^, Hnf4a^+^ proximal tubule cells proliferate less frequently (Figure 5). This suggests that expansion of proximal tubule progenitors determines the number of proximal tubule cells. Identification of proximal tubule progenitors will allow us to further investigate the developmental mechanisms of proximal tubule development.

We have previously reported that mosaic deletion of *Hnf4a* by *Six2GFPCre* in the developing mouse kidney causes a significant reduction in proximal tubule cells, phenocopying FRTS.^20^ The paucity of proximal tubules is consistent with reduced expression of proximal tubule genes, including the genes encoding glucose and phosphate transporters. However, it was unknown which genes were directly regulated by Hnf4a in proximal tubules. In this study, we performed Hnf4a ChIP-seq on newborn mouse kidneys and RNA-seq analysis of *Hnf4a* mutant kidneys by *Osr2^IresCre^.* From intersection of the ChIP-seq and RNA-seq datasets, we identified 245 Hnf4a direct target genes that were downregulated in the *Hnf4a* mutant during kidney development (Figure 7D). Among these 245 targets, the most enriched were genes associated with transmembrane transport, suggesting that the role of Hnf4a in proximal tubule development correlates with active reabsorption, the major function of the proximal tubule (Figure 7E, Table 1A). The genes associated with fatty acid metabolism were also enriched in these 245 direct target genes (Table 1B). Taking into account that proximal tubule cells are highly active in metabolism and their energy demands are primarily met by fatty acid oxidation^11,63–65^, our results suggest that Hnf4a regulates metabolic reprogramming during proximal tubule development. Consistent with this, it has been shown that Hnf4a controls lipid metabolism in Drosophila nephrocytes.^19^ Interestingly, only 3% of the Hnf4a bound genes showed differential expression in the *Hnf4a* mutant kidney. Since many proximal tubule genes are upregulated postnatally, ^67^ it is possible that Hnf4a alone is not sufficient to induce expression of the majority of its target genes and co-factors are required.

**Table 1.**
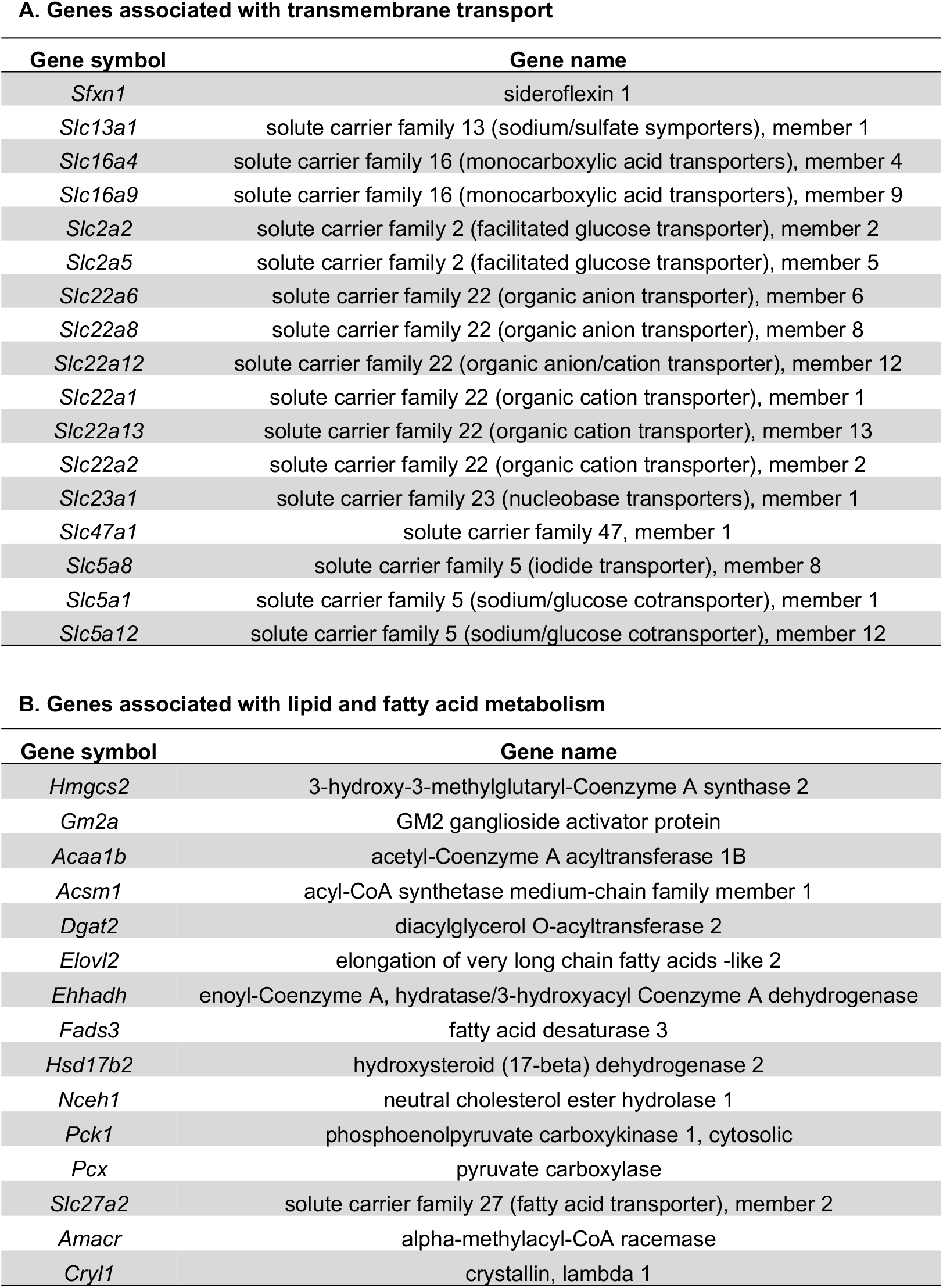
Hnf4a target genes that were downregulated in the *Hnf4a* mutant kidney

Motif analysis of genomic regions bound by a given transcription factor provides a list of other transcription factors that physically or genetically interact with the target transcription factor, sharing common target genes. Known motif analysis of our Hnf4a ChIP-seq datasets revealed that the DNA motifs for ESRRA, RXR, PPAR, and Hnf1b were enriched within the Hnf4a-bound genomic regions in the developing mouse kidney. The binding motifs of ESRRA and RXR are quite similar to the Hnf4a binding motif (Figure 6A), which could suggest that there is cooperative binding among these nuclear receptor transcription factors to activate a proximal tubule-specific transcriptional program. In contrast, similar binding motifs could imply competitive binding, since overlapping DNA motifs can lead to competition between transcription factors to activate or repress context-specific transcriptional programs.^68^ Recent studies in zebrafish have implicated retinoic acid signaling in the formation of proximal tubules, further supporting RXR as a potential co-regulator of proximal tubule development.^69, 70^ Both PPARa/g and Hnf1a/b have been implicated in proximal tubule development and function, indicating that they are good candidate co-regulators of proximal tubule development.^71–73^ PPAR transcription factors are known binding partners of RXR and one study predicted 17 common targets between Hnf4a and PPARa. *Hnf1b* is expressed in all nephron segments^74–78^ and *Hnf1b* deficiency in the nephron lineage of the mouse kidney leads to defects in nephron formation, particularly the proximal tubules, loops of Henle, and distal tubules.^78, 79^ Hnf1b is known to interact with Hnf4a and regulate common target genes.^67, 75, 80^ It has also been shown that expression of *Hnf1b* and *Hnf4a,* along with *Emx2* and *Pax8,* can convert fibroblasts into renal tubular epithelial cells, strongly suggesting that Hnf1b is a co-regulator of proximal tubule development.^81^ Further investigation is needed to elucidate the interactions among these transcription factors and their roles in proximal tubule development.

A number of protocols have been developed for *in vitro* differentaion of induced pluripotent stem cells into kidney organoids as well as transdifferentation protocols to produce renal epithelial cells for disease modeling, drug toxicity testing, and possible cell-based therapies for chronic kidney disease.^81–85^ One of the challenges of kidney organoid differentaion is achieving cellular maturity.^86^ It is known that proximal tubule cells in kidney organoids fail to undergo the maturation process.^87^ Considering our finding that Hnf4a plays a critical role in proximal tubule development, the application of Hnf4a ligands, such as linoleic acid,^88^ may be useful for the generation of mature proximal tubules in kidney organoids.

In conclusion, we examined the molecular mechanisms of *Hnf4a*-regulated proximal tubule development. We found that proximal tubule development was arrested in the absence of *Hnf4a.* The *Hnf4a* mutant kidney cannot generate mature proximal tubules. Loss of proximal tubule cells in the *Hnf4a* mutant mice caused postnatal lethality, highlighting the importance of functional proximal tubules for survival. In the *Hnf4a* mutant kidney, there is an increase in Cdh6^high^, LTL^low^ presumptive proximal tubule cells. We definitively showed that Cdh6^+^ cells in the developing kidney are proximal tubule progenitors. These results suggest that *Hnf4a* is required for proximal tubule progenitors to differentiate into mature proximal tubule cells. Genome-wide analysis of Hnf4a binding sites in the kidney and transcriptomic analysis of the *Hnf4a* mutant kidney indicate that Hnf4a directly regulates expression of multiple genes involved in transmembrane transport and metabolic processes in the proximal tubule.

## Author contributions

S.S.M. performed mouse experiments. S.S.M. and E.C. performed ChIP-seq. E.C. performed RNA-seq. S.S.M. and J.P. designed the experiments, analyzed the data, and cowrote the manuscript. S.S.M made the figures. All authors approved the final version of the manuscript.

## Acknowledgments

The authors thank Steve Potter and Mike Adam for help with single cell RNA-seq data. We also thank the Confocal Imaging Core (CIC) and the DNA Sequencing and Genotyping Core (DSGC) at CCHMC. This work was supported by the National Institute of Diabetes and Digestive and Kidney Diseases, National Institutes of Health F31 DK120164 to S.S.M. and R01 DK120847 to J.P.

## Supplemental Material

Supplemental Figure 1. Deletion of *Hnf4a* does not affect the formation of the S-shaped body.

Supplemental Figure 2. Deletion of *Hnf4a* does not affect the formation of other nephron segments in the newborn kidney.

Supplemental Figure 3. Mature proximal tubule markers are absent in the *Hnf4a* mutant kidney.

Supplemental Figure 4. The glomerulus is connected to Cdh6+ tubules in the Hnf4a mutant and control kidneys.

Supplemental Figure 5. Deletion of *Hnf4a* does not affect the formation of other nephron segments in the postnatal kidney.

Supplemental Figure 6. *Cdh6* expression persists postnatally in the *Hnf4a* mutant and control kidneys.

Supplemental Figure 7. Expression of top marker genes that are enriched in early and mature proximal tubules.

Supplemental Table 1. Genome-wide mapping of Hnf4a binding sites in the mouse kidney at P0 (ChIP-seq)

Supplemental Table 2. Differential gene analysis of the Hnf4a mutant kidney at P0 (RNA-seq)

Supplemental Table 3. Intersection of ChIP-seq and RNA-seq

