## supplemental figures for "*Hnf4a* is required for the development of Cdh6-expressing progenitors into proximal tubules in the mouse kidney"

Short title: Hnf4a in proximal tubule development

Corresponding author:

Joo-Seop Park

Cincinnati Children's Hospital Medical Center

Location R1566, ML7007

3333 Burnet Avenue

Cincinnati, OH 45229

**Supplemental Figure 1.** Deletion of *Hnf4a* does not affect the formation of the S-shaped body. Representative image of *n*=3. Stage E18.5. Scale bar, 25µm.

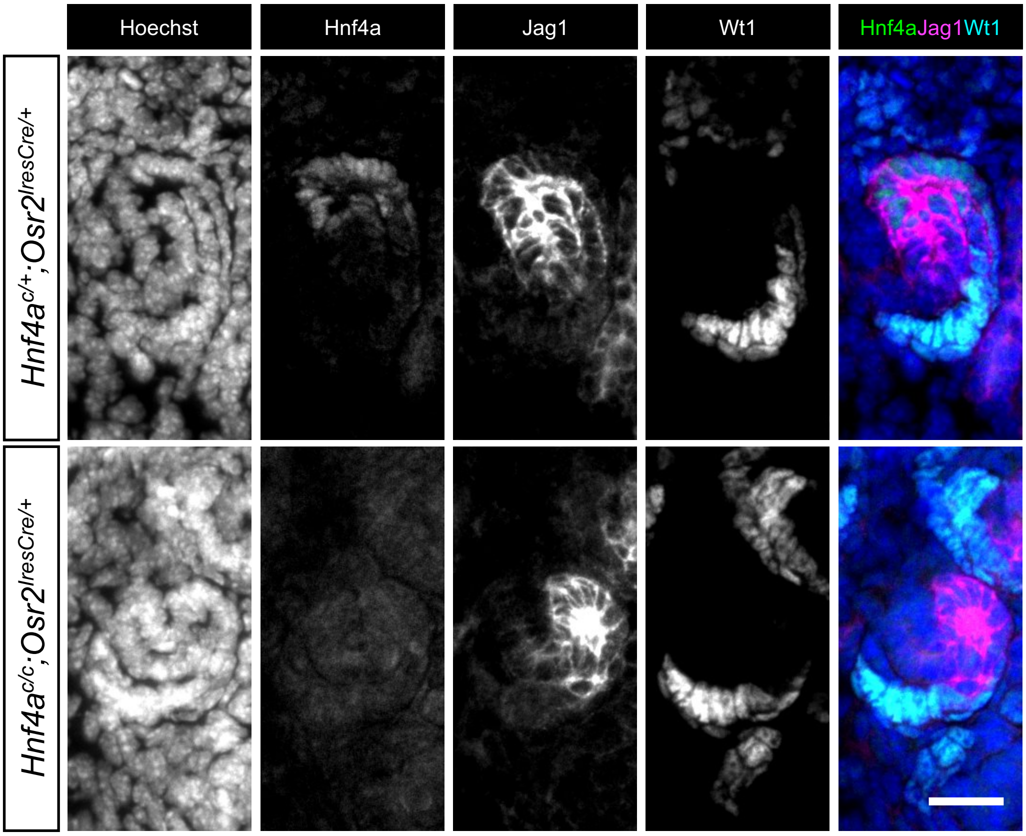

**Supplemental Figure 2.** Deletion of *Hnf4a* does not affect the formation of other nephron segments in the newborn kidney. (A) Immunofluorescence staining on podocytes (Wt1) , loops of Henle (Slc12a1), and distal tubules (Slc12a3) shows no difference between mutant and control. Representative image of *n*=3. Stage P0. Scale bar, 100µm. (B) RNA-seq analysis of *Hnf4a* mutant kidneys at P0 showed a significant decrease in expression of PT genes but not podocyte (Pod), loop of Henle (LOH), nor distal tubule (DT) genes.

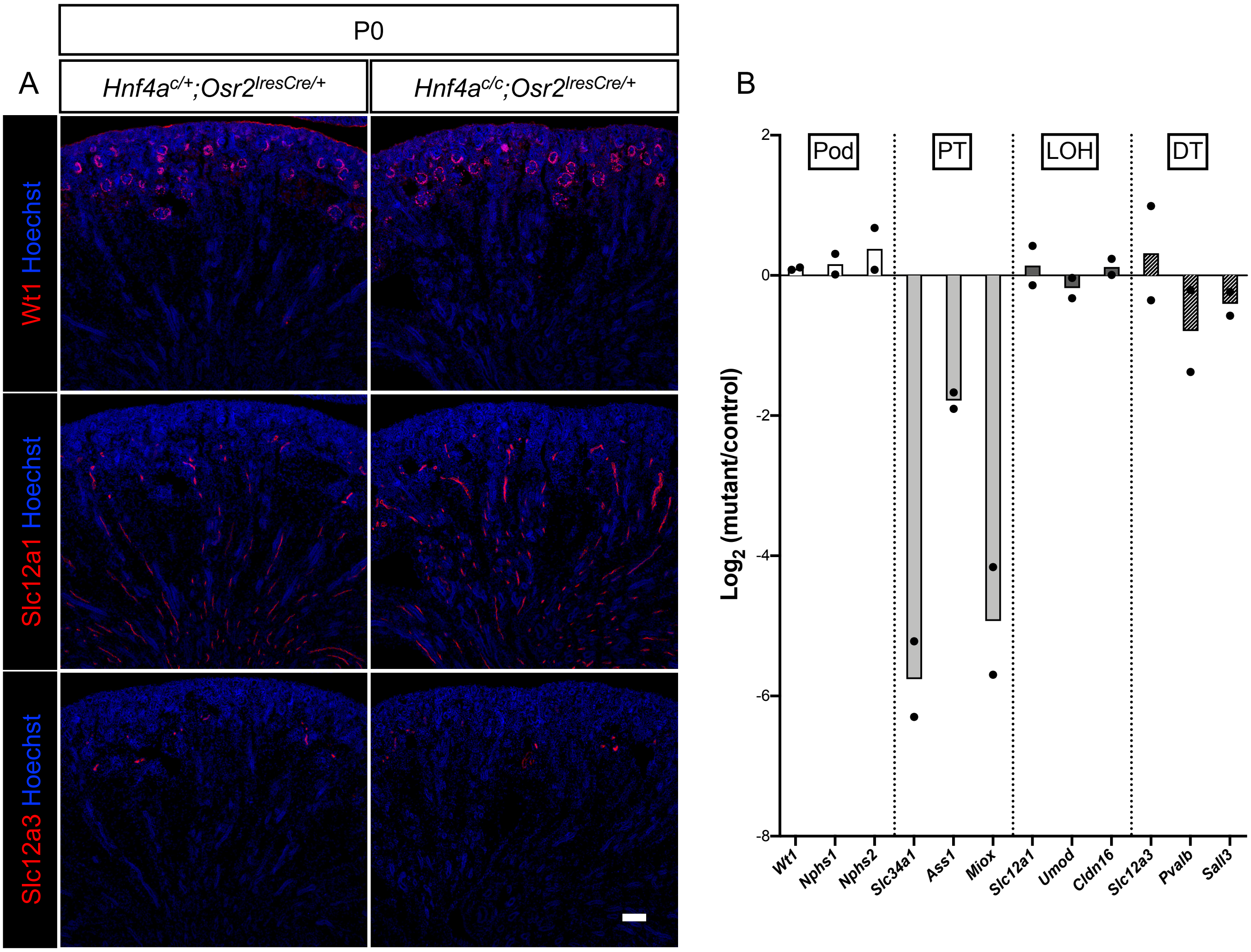

**Supplemental Figure 3.** Mature proximal tubule markers are absent in the *Hnf4a* mutant kidney. Representative image of *n*=3. Stage P0. Scale bar, 100µm.

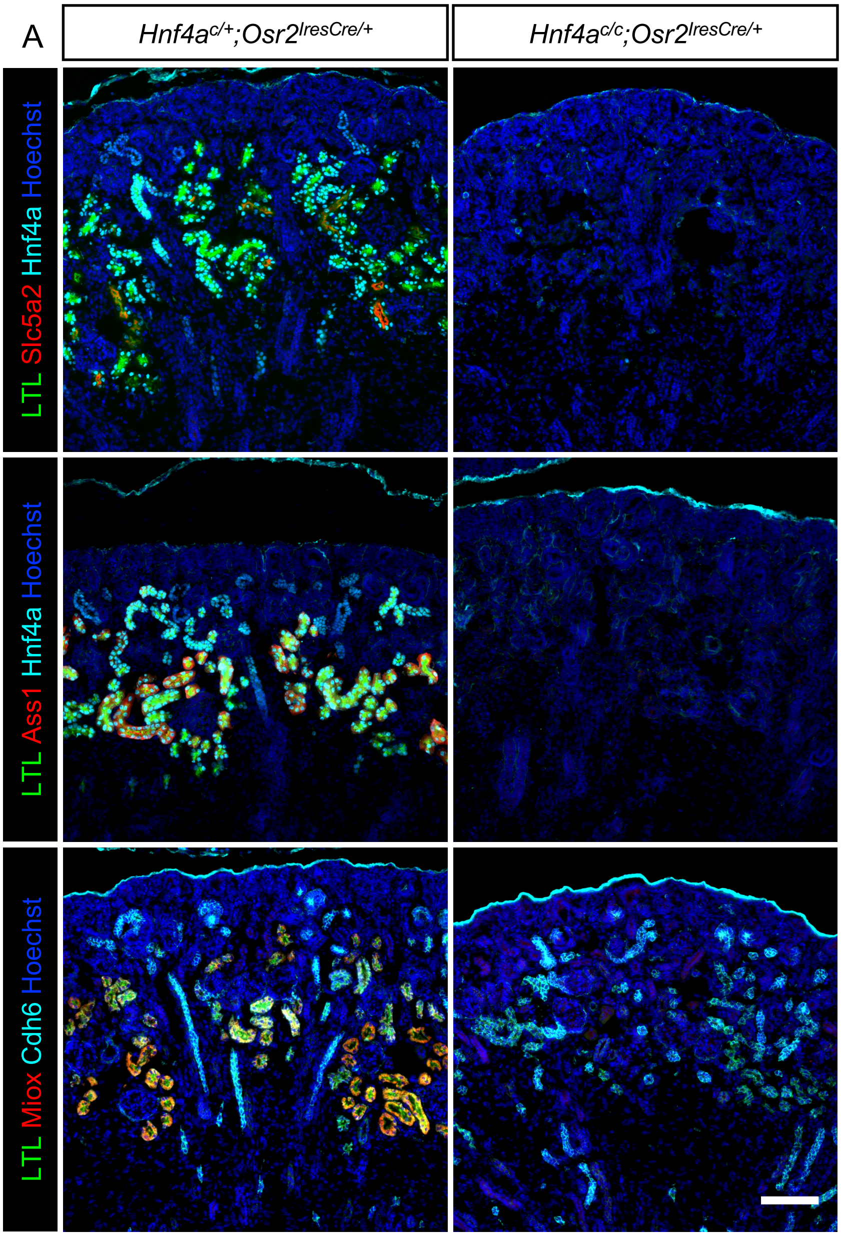

**Supplemental Figure 4.** The glomerulus is connected to Cdh6$^{+}$ tubules in the *Hnf4a* mutant and control kidneys. Representative image of *n*=3. Stage E18.5. Scale bar, 25µm.

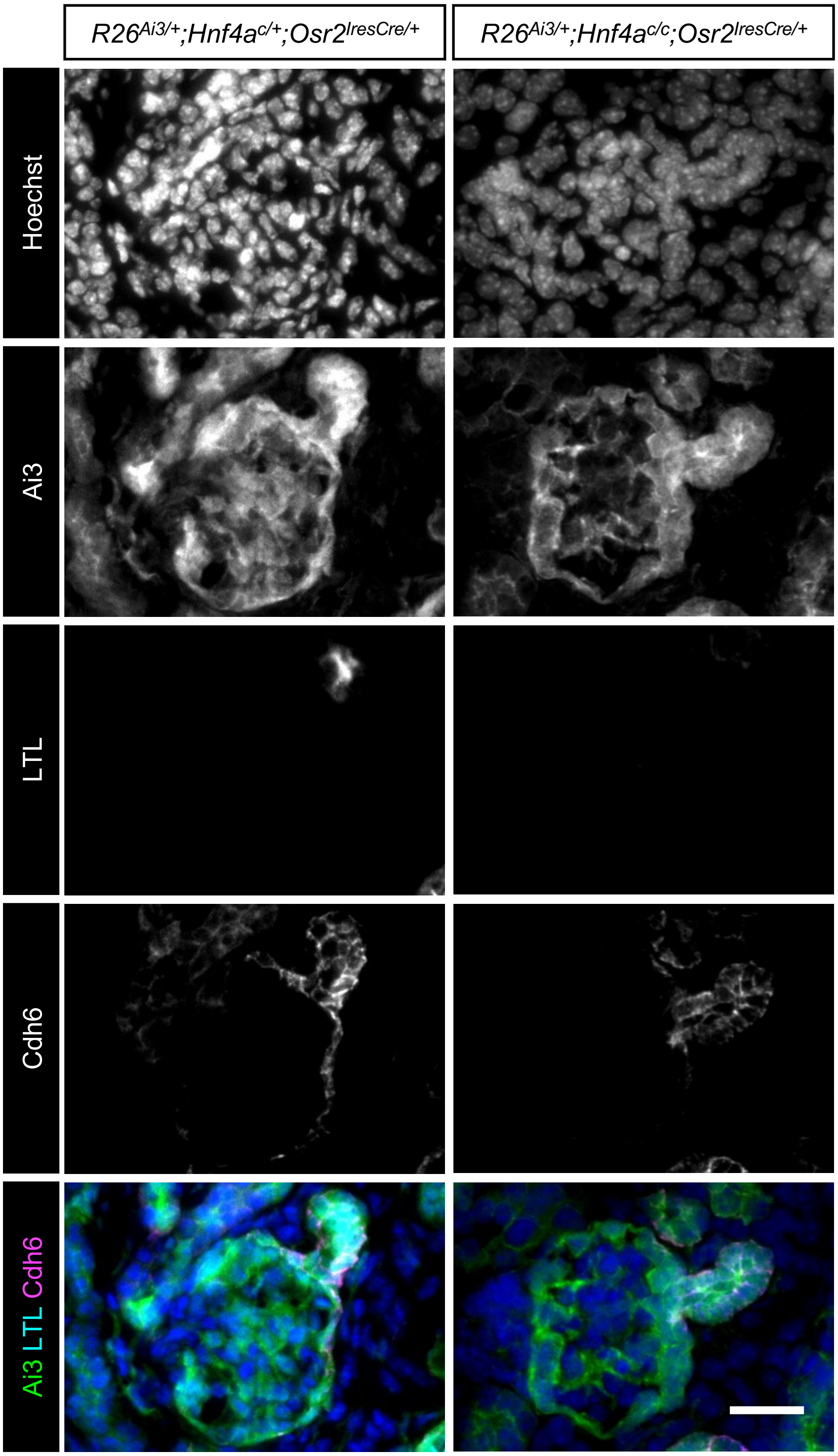

**Supplemental Figure 5.** Deletion of *Hnf4a* did not affect the formation of other nephron segments in the postnatal kidney. Wt1, Slc12a1,a and Slc12a3 mark podocytes, loops of Henle, and distal tubules, respectively. Representative image of *n*=3. Stage P7. Scale bar, 100μm.

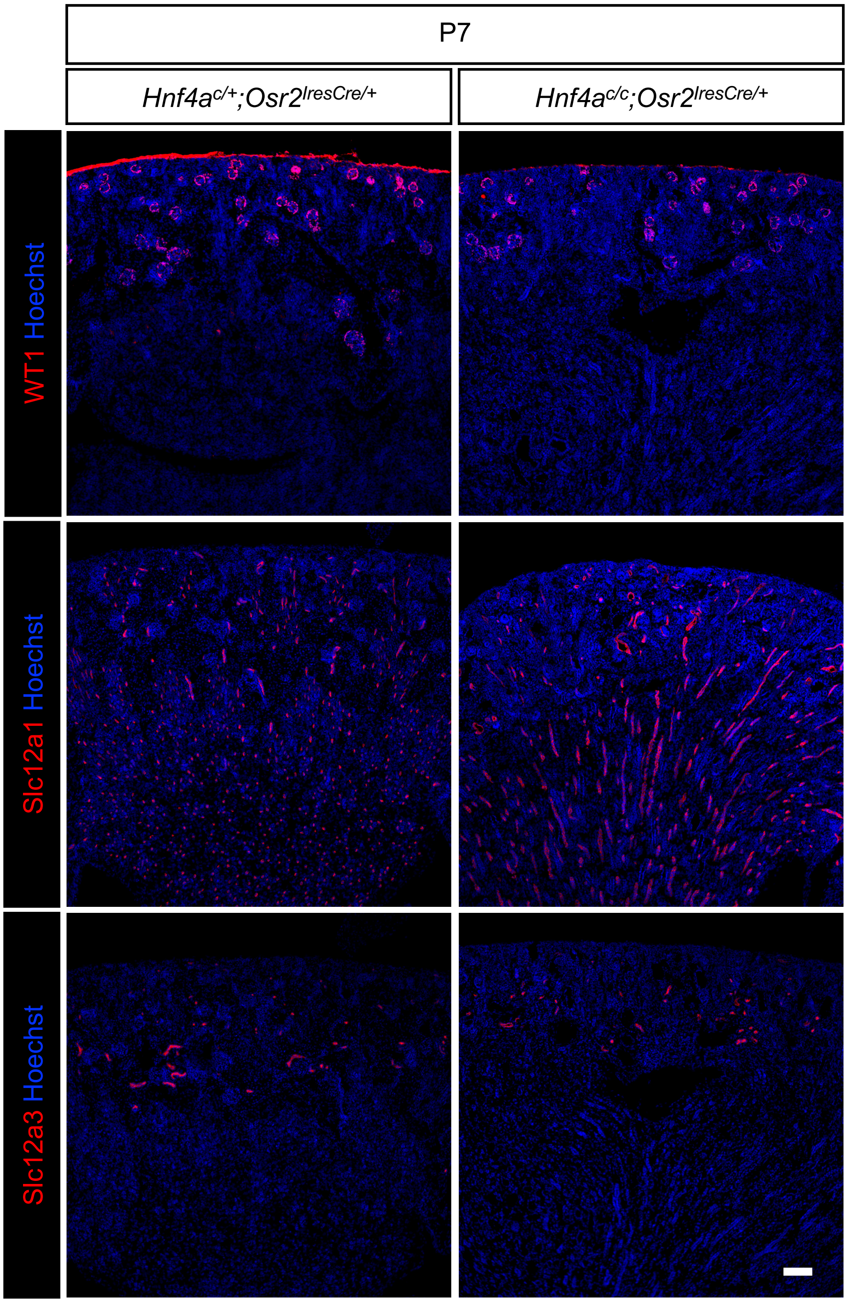

**Supplemental Figure 6.** *Cdh6* expression persists postnatally in the *Hnf4a* mutant and control kidneys. Representative image of *n*=3. Scale bar, 100μm.

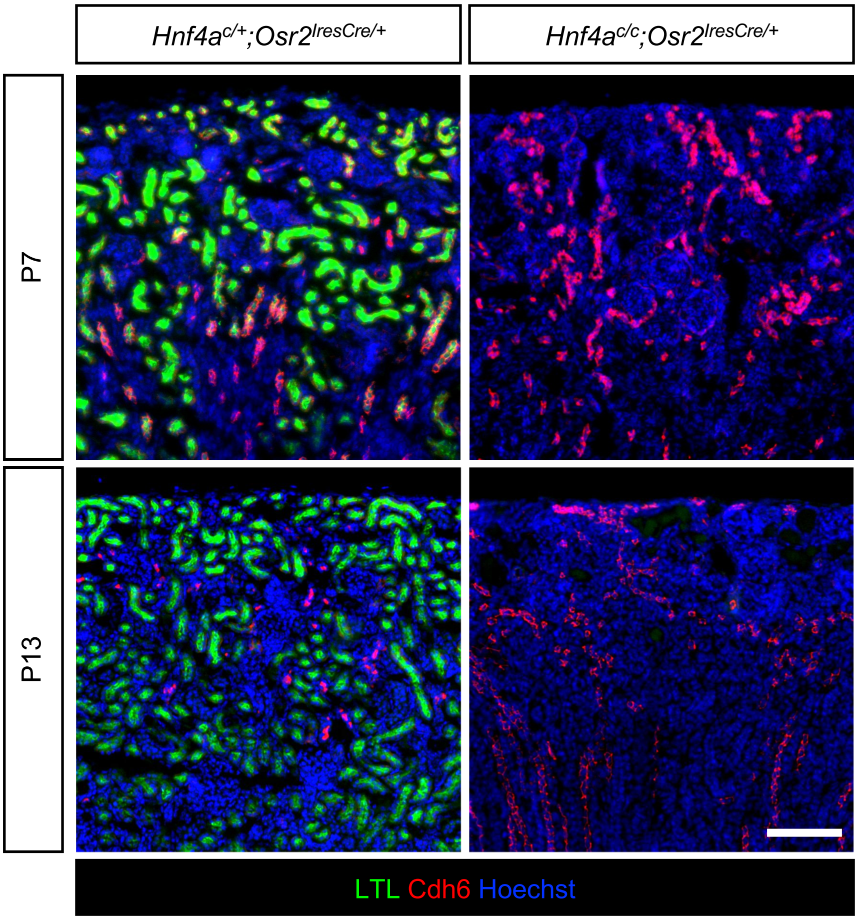

**Supplemental Figure 7.** Expression of top marker genes that are enriched in early and mature proximal tubules. These genes were previously identified from single cell RNA-seq data of the newborn mouse kidney (ref 8). The genes enriched in early proximal tubules (blue) were not affected in the *Hnf4a* mutant kidney. The genes enriched in mature proximal tubules (red) were downregulated in the *Hnf4a* mutant kidney. Dotted lines represent y= $\pm \log_2(1.5)$. The median of the data is represented as a straight black line. Stage P0.

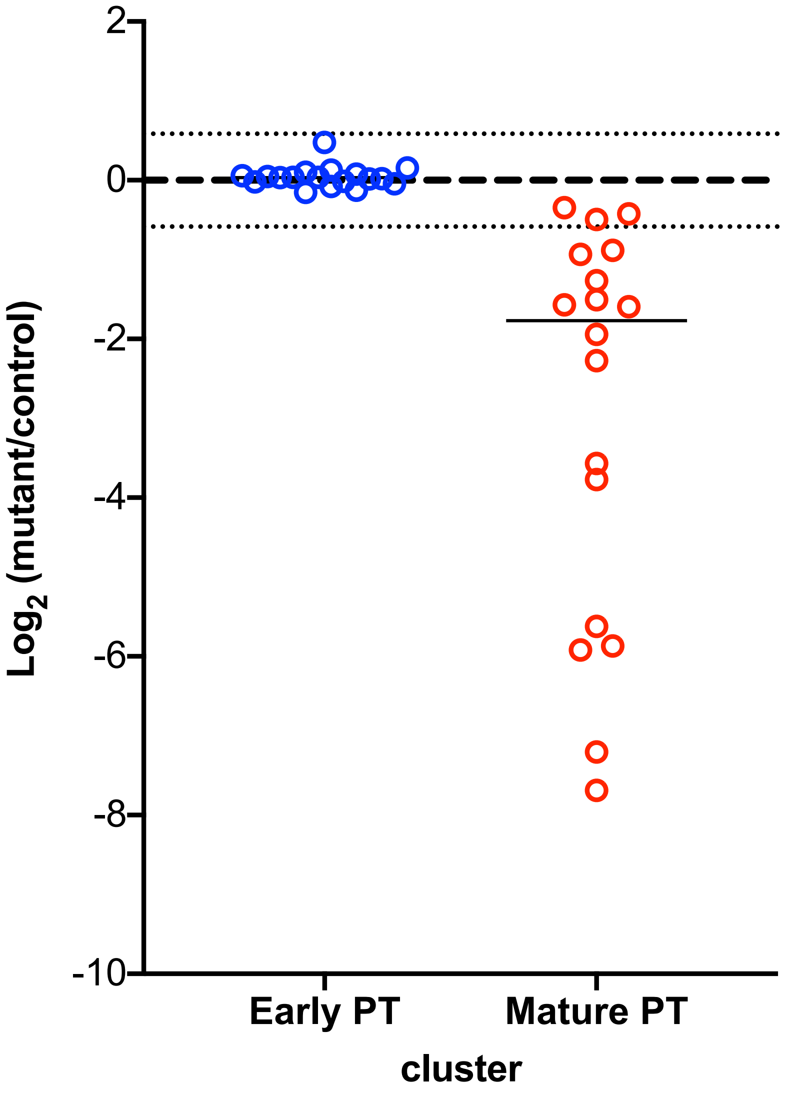
